# Dynamic Brain Network Changes Associated with Successful Smoking Cessation

**DOI:** 10.1101/2025.09.13.676003

**Authors:** Jooyeon Jamie Im, Hyeonjin Kim, Jeung-Hyun Lee, Hyung Jun Park, Hee-Kyung Joh, Woo-Young Ahn

**Affiliations:** Department of Psychology, Seoul National University, Seoul, Republic of Korea; Center for Health Promotion, Samsung Medical Center, Seoul, Republic of Korea; Department of Clinical Medical Sciences, College of Medicine, Seoul National University, Seoul, Republic of Korea; Department of Family Medicine, Seoul National University Health Service Center, Seoul, Republic of Korea; Department of Medicine, Seoul National University College of Medicine, Seoul, Republic of Korea; Department of Family Medicine, Seoul National University Hospital, Seoul, Republic of Korea; Department of Brain and Cognitive Sciences, Seoul National University, Seoul, Republic of Korea; AI Institute, Seoul National University, Seoul, Republic of Korea

**Keywords:** Smoking cessation, Resting-state functional magnetic resonance imaging, Salience network, Executive network, Default mode network

## Abstract

**Background:** Tobacco smoking continues to be a leading cause of preventable morbidity and mortality globally, with the success rate of unaided cessation remaining consistently low. Understanding the neurobiological mechanisms of smoking cessation is crucial for improving quit rates. However, there has been a lack of studies examining brain network changes associated with smoking cessation over time. In this study, we aimed to investigate longitudinal changes in the functional connectivity (FC) of large-scale brain networks underlying smoking cessation outcomes using resting-state functional magnetic resonance imaging (fMRI).

**Methods:** A total of 98 treatment-seeking smokers participated in a 6-week cessation program and underwent resting-state fMRI scans before and after the intervention. Independent component analysis identified the salience network (SN), executive control network (ECN), and default mode network (DMN) components, and region of interest (ROI)-to-ROI FC was compared between successful and unsuccessful quitters using a group × time mixed-effects model. Correlations with smoking-related measures were explored.

**Results:** Significant group-by-time interaction effects were found in FC, particularly involving connections between SN and ECN, as well as between the SN and DMN. Specifically, successful quitters exhibited greater baseline FC in the SN-ECN and SN-DMN circuits, which tended to normalize during the cessation process. Exploratory correlational analyses revealed trends suggesting that stronger pre-quit connectivity between the SN and ECN was associated with greater withdrawal severity and longer smoking history in successful quitters.

**Conclusions:** Taken together, the normalization of initially elevated pre-quit FC in SN-ECN and SN-DMN circuits may reflect an adaptive neural process that supports successful withdrawal management and attentional reallocation during cessation. The identification of these neural substrates not only enhances our mechanistic understanding of smoking cessation over time but also underscores the need for targeted interventions that focus on these neural circuits to enhance quit outcomes.

## 1. Introduction

Tobacco smoking remains a leading cause of preventable morbidity and mortality globally, posing a major public health concern (1). Approximately 70% of smokers express a desire to quit, yet the success rate of unaided cessation hovers below 10%, highlighting a stark disconnect between motivation and the capacity to overcome nicotine dependence (2). This disconnect underscores the critical need to investigate the underlying neurobiological mechanisms of smoking cessation to develop more effective interventions that can improve quit rates.

Recently, there has been growing interest in understanding the functional connectivity (FC) of large-scale brain networks using resting-state functional magnetic resonance imaging (fMRI). Notably, research has primarily focused on three key networks including the salience network (SN), executive control network (ECN), and default mode network (DMN), as these are considered the most clinically relevant (3–6). The SN, which includes regions such as the anterior insula and anterior cingulate cortex, is involved in detecting and responding to important stimuli. The ECN, also known as the frontoparietal network, comprises areas such as the dorsolateral prefrontal cortex and posterior parietal cortex, and is implicated in cognitive control, decision-making, and goal-directed behavior. Lastly, the DMN, including regions such as the medial prefrontal cortex, posterior cingulate cortex, and lateral parietal cortex, has been associated with self-referential processing and mind-wandering.

Most previous studies examining large-scale brain networks in smokers have either compared smokers to non-smokers or investigated the effects of abstinent versus satiated states within smokers, but few have specifically addressed smoking cessation outcomes. For example, studies comparing smokers and non-smokers often report decreased FC, notably in the SN (7–9), ECN (9–12), and DMN (12, 13), among smokers. Additionally, studies comparing abstinent and satiated states in smokers have shown that FC patterns within and between brain networks vary depending on the smoking state (9, 14, 15). In particular, decreased FC between the SN and ECN, along with increased FC between the SN and DMN, has been identified as a neural signature of smokers during acute abstinence. While these studies have provided valuable insights into the effects of smoking and the impact of nicotine abstinence on brain networks, the relationship between these connectivity alterations and smoking cessation outcomes remains unclear.

Among the limited number of studies examining the association between FC and smoking cessation outcomes, most have relied on a seed-based approach in resting-state fMRI analyses, rather than utilizing network-based methods. These studies have focused on regions such as the insula and striatum, and notable findings highlight the involvement of insula-centered circuits and their connectivity changes related to withdrawal and cessation outcomes (16–19). While useful for examining regional connectivity changes, seed-based methods may not fully capture the complex interactions across the entire brain network (20). Furthermore, most of the studies investigating resting-state FC with smoking cessation outcomes have only used baseline scans collected before quit attempts to predict later smoking outcomes, limiting the understanding of longitudinal changes in FC.

Taken together, there is a gap in research that has explored the longitudinal changes in large-scale brain network connectivity involved in smoking cessation. Here, the present study aims to address this gap by examining these changes in smokers during a quit attempt. Specifically, we compared smokers who successfully maintained abstinence with those who relapsed during a 6-week cessation program. We hypothesized that successful quitters would exhibit distinct patterns of FC in the SN, ECN, and DMN compared to smokers who relapsed, with successful quitters showing stronger connectivity between the SN and ECN, and weaker connectivity between the SN and DMN at baseline, and these differences may stabilize or diminish as the brain adapts during the cessation process.

## 2. Methods and Materials

### 2.1. Participants and Procedures

Smokers who reported an intention to quit smoking were recruited through flyers on the university campus and online postings on local community websites. The inclusion criteria for the study were as follows: smoking a minimum of five cigarettes per day for the past year, or an equivalent amount of e-cigarettes as defined by at least five sessions of 15 puffs each or 10 minutes of smoking at each session (21), and willing to make a quit attempt and enroll in the smoking cessation program at a university health center. Exclusion criteria included major psychiatric or medical illness, use of psychoactive medications, other substance abuse, including alcohol, as assessed by the Structured Clinical Interview for DSM-5 Disorders (22), use of illicit drugs based on a urine drug screen (One-Step Drug Abuse Test Dip Card from WHPM, Inc.), and contraindications for magnetic resonance imaging (MRI).

The study protocol involved three laboratory visits: a screening visit, a pre-quit MRI scan visit preceding the smoking cessation program, and a post-quit MRI scan visit upon its completion. Prior to both MRI scans, participants were asked to abstain from smoking, alcohol use, and caffeine consumption for at least 8 hours. The abstinence was verified with exhaled carbon monoxide (CO) levels using a Micro Smokerlyzer (Bedfont Scientific Ltd.) and alcohol concentration level using a breathalyzer (AL8800 ALCOSCAN, Sentech Korea). Participants were expected to have CO levels lower than 12 ppm after 8 hours of abstinence.

The smoking cessation program, lasting 5 to 6 weeks, included weekly visits to the clinic and offered both pharmacological and psychosocial treatments. Consistent with previous studies, smoking abstinence was defined as continuous abstinence over the last 4 weeks of the program (23, 24). Participants who remained abstinent during this period were classified as successful quitters, and those who did not maintain abstinence as unsuccessful quitters, based on self-reports and cross-validation by CO levels of 5 ppm or below to be confirmed as successful quitters (25). The study procedures were approved by the Institutional Review Boards of Seoul National University and were performed in accordance with the ethical standards of the Declaration of Helsinki. Written consent form was obtained from all participants to confirm their voluntary participation.

### 2.2. Clinical Assessments

Demographic information and self-report measures including smoking-related assessments were obtained at the screening visit after confirming eligibility. Nicotine dependence was assessed with the Fagerström Test of Nicotine Dependence (FTND) (26, 27) and Cigarette Dependence Scale (CDS-12) (28). Nicotine craving was evaluated with the Brief Questionnaire on Smoking Urges (QSU-Brief) (29). Nicotine withdrawal was assessed with the Cigarette Withdrawal Scale (CWS) (30) and Minnesota Nicotine Withdrawal Scale (MNWS) (31, 32). Daily cigarette consumption was initially recorded as a categorical variable as follows: 0 for none, 1 for 1–4 cigarettes, 2 for 5–10 cigarettes, 3 for 11–20 cigarettes, and 4 for greater than 20 cigarettes per day. During the latter two-thirds of the study period, we began collecting the daily cigarette consumption data as a continuous variable, detailing the exact number of cigarettes smoked per day. For the 20% of our sample who had only provided categorical responses, we inferred daily averages by assigning median values to each category: 0 equating to 0 cigarettes; 1 to 3 cigarettes (median of 1 and 4); 2 to 7.5 cigarettes (median of 5 and 10); 3 to 15.5 cigarettes (median of 11 and 20); and 4 to 30 cigarettes (median of 20 and 40). For electronic cigarettes, 15 puffs or 10 minutes of smoking were considered equivalent to one session, comparable to one regular cigarette (21). Duration of smoking was calculated by subtracting the age of smoking initiation from the current age.

### 2.3. MRI Data Acquisition and Preprocessing

All MRI scanning was conducted on a Siemens TIM Trio 3T scanner using a 32-channel head coil. A high-resolution T1-weighted structural images were acquired using MPRAGE sequence (TR = 2300 ms, TE = 2.36 ms, field of view = 256 mm^2^, flip angle=9°, slices = 224, and slice thickness = 1 mm). Resting-state fMRI was performed using an T2*-weighted, single-shot echo, echo-planar imaging sequence (TR = 1500 ms, TE = 30 ms, field of view = 256 mm^2^, flip angle = 85°, slices = 64, slice thickness = 2.3 mm, and 250 volumes for a total duration of 6 min 44 s). During the resting-state fMRI scan, participants were instructed to rest with their eyes open while fixating on a cross in the center of the screen. An infrared camera attached to the head coil was used to monitor alertness. These images were collected as part of a 60-min scanning session in the following order: a localizer, T1-weighted structural MRI, resting-state fMRI, task-based fMRI, and neuromelanin-sensitive MRI.

### 2.4. Data Preprocessing and Analysis

Resting-state fMRI analyses consisted of 1) preprocessing, 2) independent component analysis (ICA), 3) first-level calculation of resting-state FC matrices of the ROIs based on the ICA results, 4) second-level group (successful quit group vs unsuccessful quit group) x time (pre and post) interaction analyses controlling for age, gender, and smoking duration, 5) and correlation analyses with self-report measures. Preprocessing and analyses of the resting-sate fMRI data were performed using the CONN toolbox (RRID:SCR_009550) release 21.a (33, 34) with SPM12.

#### Preprocessing

Functional and anatomical data were preprocessed using a standard preprocessing pipeline including realignment, slice timing correction, outlier detection, direct segmentation, MNI-space normalization, and smoothing. Functional data alignment was performed using the SPM realign & unwarp procedure, with all scans coregistered to the first scan of the initial session via a least squares method and a 6-parameter (rigid body) transformation. B-spline interpolation was applied to correct for motion and magnetic susceptibility interactions, and temporal misalignment between different slices of the functional data (acquired in interleaved Siemens order) was corrected following SPM slice-timing correction procedure. Potential outlier scans were identified using ART as acquisitions with framewise displacement above 0.5 mm or global BOLD signal changes above 3 standard deviations, and a reference BOLD image was computed for each subject by averaging all scans excluding outliers. Functional and anatomical data were then normalized into standard MNI space, segmented into grey matter, white matter, and CSF tissue classes, and resampled to 2 mm isotropic voxels. Lastly, functional data were smoothed using spatial convolution with a Gaussian kernel of 8 mm full width half maximum (FWHM). In addition, functional data were denoised using a standard denoising pipeline, which involved regressing out potential confounding effects. These included white matter timeseries (5 CompCor noise components), CSF timeseries (5 CompCor noise components), motion parameters and their first order derivatives (12 factors), outlier scans (below 58 factors), session and task effects and their first order derivatives (4 factors), and linear trends (2 factors) within each functional run. The BOLD timeseries was then bandpass filtered between 0.008 Hz and 0.09 Hz.

#### Group Independent Component Analysis

Group-level independent component analyses (group-ICA) were performed to estimate 20 temporally coherent networks from the resting-state fMRI data across all participants and sessions (35). The BOLD signal from every timepoint and voxel in the brain was concatenated across participants and sessions along the temporal dimension. A singular value decomposition of the z-score normalized BOLD signal with 64 components separately for each participant and session was used as a subject-specific dimensionality reduction step. The dimensionality of the concatenated data was further reduced using a group-level singular value decomposition with 20 components, and a fast-ICA fixed-point algorithm with hyperbolic tangent (G1) contrast function was used to identify spatially independent group-level networks from the resulting components. Lastly, GICA3 back-projection was used to compute ICA maps associated with these same networks separately for each participant and session. Each ICA map was visually checked for the SN, ECN, and DMN networks (3) and cross-validated with the external template provided in the CONN toolbox. The ICA maps were thresholded at Z ≥ 3.0 for the group-level connectivity analysis.

#### First-Level and Second-Level Network Analyses

All clusters within the SN, ECN, DMN maps were selected as the region of interests (ROIs). In the first level analysis, ROI-to-ROI connectivity matrices were estimated characterizing the FC connectivity between each pair of regions among 21 ROIs, resulting in a total of 210 (=21*(21–1)/2) connections. FC connectivity strength was represented by Fisher-transformed bivariate correlation coefficients, characterizing the association between their BOLD signal timeseries. A linear mixed-effects model was conducted to assess the group (successful quit group and unsuccessful quit group) x time (pre and post) interaction effects controlling for age, gender, and smoking duration.

Significance was determined at the level of clusters/groups of connections. Connection-level hypotheses were evaluated using multivariate parametric statistics with random-effects across participants and sample covariance estimation across multiple measurements. Inferences were performed at the level of individual clusters (groups of similar connections). Cluster-level inferences were based on parametric statistics within- and between-each pair of networks (functional network connectivity), with networks identified using a complete-linkage hierarchical clustering procedure based on ROI-to-ROI anatomical proximity and functional similarity metrics. Results were corrected across multiple comparisons using FDR across the entire set of possible pairwise clusters. The statistical significance level was set at connection-level threshold of p < 0.05 (uncorrected) and cluster-level threshold of p-FDR < 0.05.

#### Correlational Analyses between Network-Based Resting-State FC and Self-Report Measures

Statistical analyses of the demographic and self-report data were conducted using R software (version 4.3.2). Baseline characteristics of groups were compared using independent t-test for continuous variables, and chi-square test for categorical variables. Subsequently, we performed correlational analysis to explore the relationships between FC patterns and self-report measures including smoking-related variables. To do this, we extracted FC values from the significant clusters of the baseline and follow-up scans for each participant and analyzed for correlations with smoking-related variables using Pearson’s correlation coefficients.

## 3. Results

### Demographic and Smoking-Related Characteristics

Of 98 participants who had both pre- and post-quit resting-state fMRI scans, 6 participants were excluded due to using different MRI acquisition parameters, and 1 participant was excluded for prematurely attempting to quit before the smoking cessation program started. Consequently, 91 participants were included in the final analysis. Demographic and smoking-related characteristics for the unsuccessful and successful quit groups are summarized in **Table 1**. According to the smoking cessation criterion defined in our study, which is complete abstinence during the final 4 weeks, 26 participants were classified as successful quitters, and 65 participants were classified as unsuccessful quitters. At the pre-quit visit, successful quitters did not differ significantly from unsuccessful quitters in terms of age, sex, education, daily smoking amount, CO level, or the severity of nicotine dependence as measured by the FTND scores. Successful quitters had a longer smoking duration (p = 0.006) than unsuccessful quitters at the pre-quit visit. At the post-quit visit, successful quitters had higher program participation rates (p = 0.003) and lower CO levels (p = 0.001) than unsuccessful quitters.

**Table 1.** Demographic and Smoking-Related Characteristics.

| Characteristic | Unsuccessful quitters<br>(n = 65) | Successful quitters<br>(n = 26) | Test<br>Statistic | *p*-value |
| --- | --- | --- | --- | --- |
| Age, years | 25.35 ± 3.99 | 26.23 ± 3.37 | -0.97 | 0.3 |
| Sex, male:female | 55:10 | 22:4 | 0.00 | 1.0 |
| Education, year | 16.12 ± 2.25 | 16.54 ± 2.93 | -0.73 | 0.5 |
| Smoking amount, cigarette/day | 11.67 ± 6.92 | 9.50 ± 4.01 | 1.5 | 0.14 |
| Smoking duration, year | 4.57 ± 4.15 | 7.31 ± 4.25 | -2.8 | 0.006** |
| FTND score | 2.54 ± 1.75 | 1.92 ± 1.55 | 1.6 | 0.12 |
| Program participation, % | 53.08 ± 31.99 | 74.36 ± 22.72 | -3.1 | 0.003** |
| CO level at pre-quit, ppm | 4.80 ± 4.60 | 4.00 ± 4.14 | 0.77 | 0.4 |
| CO level at post-quit, ppm | 4.46 ± 3.86 | 1.85 ± 1.05 | 3.4 | 0.001** |

Data are presented as mean ± standard deviation, unless otherwise indicated.
\*\**p*<0.01; \**p*<0.05.
Abbreviations: CO: Carbon Monoxide; FTND: Fagerström Test of Nicotine Dependence.

### Group ICA

Of the 20 group-ICA components, 5 components were identified as the SN, ECN, and DMN:

1. **the SN**, including the bilateral superior frontal gyrus, middle frontal gyrus, dorsal anterior cingulate cortex (ACC)/paracingulate gyrus, anterior insula, right cerebellum, and left precuneus (**Supplementary Figure 1A**),
2. **the left ECN**, including the left dorsolateral prefrontal cortex (DLPFC) and angular gyrus/inferior parietal gyrus (**Supplementary Figure 1B**) and **right ECN**, including the right DLPFC, angular gyrus/inferior parietal gyrus, orbital frontal gyrus, inferior temporal gyrus, and cerebellum (**Supplementary Figure 1C**),
3. **the DMN**, including the left medial PFC, angular gyrus, and inferior temporal gyrus (**Supplementary Figure 1D**), and PCC (**Supplementary Figure 1E**).

Spatial cross-correlation of these networks with the external template revealed a mean r of 0.34 with a range between 0.20 and 0.44.

### Network Analysis

As shown in **Figure 1**, significant group-by-time interaction effects on FC were found across 11 inter-network connections, consisting of 5 SN-DMN and 6 SN-EN connections (F(3, 84) = 5.54, p-uncorrected = 0.0016, p-FDR = 0.0097). Details about the peak MNI coordinates and their corresponding anatomical regions for each significant connection are summarized in **Table 2**. No significant group differences were observed at either the pre- or post-quit visits.

**Figure 1.**
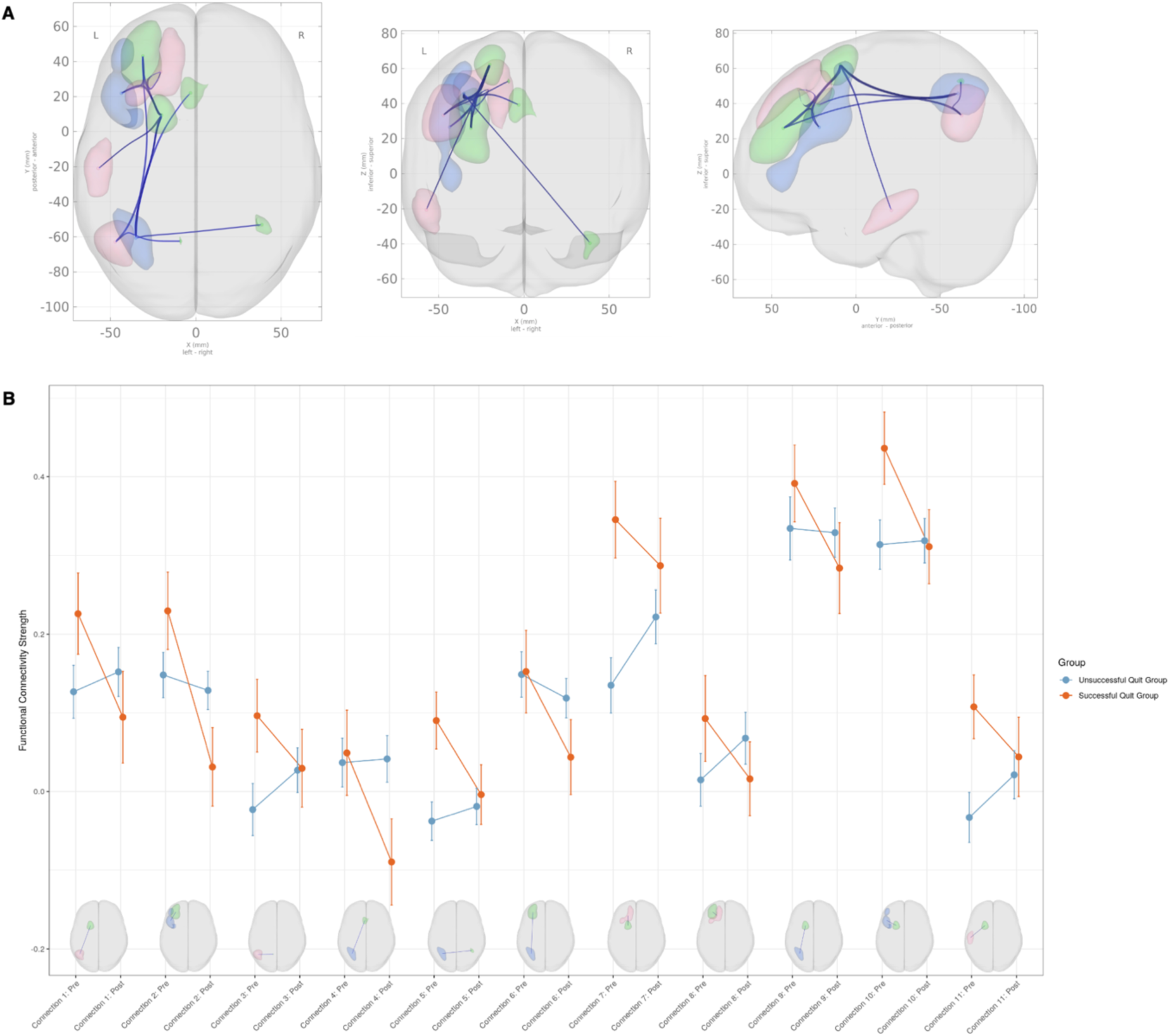
(A) Significant group (successful quit group x unsuccessful quit group) by time (pre x post) interaction. (B) Interaction plots. The clusters belong to the Salience Network (green), Default Mode Network (red), and Executive Control Network (blue). Please refer to Table 2 for detailed information on the anatomical regions.

**Table 2.** Regions with significant group (successful quit group x unsuccessful quit group) by time (pre x post) interaction on functional connectivity.

| Conne-<br>ction | Network | Anatomical region | Peak MNI coordinates |  | T value |
| --- | --- | --- | --- | --- | --- |
| 1 | SN – DMN | L. SFG – L. Angular gyrus | -20, 9, 62 | -47, -63, 33 | -2.83 |
| 2 | SN – ECN | L. MFG – L. DLPFC | -31, 43, 25 | -44, 22, 27 | -2.74 |
| 3 | SN – DMN | L. Precuneus – L. Angular gyrus | -9, -62, 53 | -47, -63, 33 | -2.45 |
| 4 | SN – ECN | L. ACC – L. IPG | -3, 22, 39 | -35, -61, 46 | -2.54 |
| 5 | SN – ECN | R. Cerebellum – L. IPG | 39, -53, -40 | -35, -61, 46 | -2.26 |
| 6 | SN – ECN | L. MFG – L. IPG | -31, 43, 26 | -35, -61, 46 | -2.14 |
| 7 | SN – DMN | L. SFG – L. SFG | -20, 9, 62 | -20, 34, 48 | -2.44 |
| 8 | SN – DMN | L. MFG – L. SFG | -31, 43, 26 | -20, 34, 48 | -2.29 |
| 9 | SN – ECN | L. SFG – L. IPG | -20, 9, 62 | -35, -61, 46 | -1.99 |
| 10 | SN – ECN | L. SFG – L. DLPFC | -20, 9, 62 | -44, 22, 27 | -2.07 |
| 11 | SN – DMN | L. SFG – L. ITG | -20, 9, 62 | -57, -21, -21 | -2.07 |

Abbreviations: ACC: Anterior Cingulate Cortex; DLPFC: Dorsolateral Prefrontal Cortex; DMN: Default Mode Network; ECN: Executive Control Network; IPG: Inferior Parietal Gyrus; ITG: Inferior Temporal Gyrus; L: Left; MFG: Middle Frontal Gyrus; MNI: Montreal Neurological Institute; R: Right; SN: Salience Network; SFG: Superior Frontal Gyrus.

Among the SN-DMN connections, the successful smoking cessation group generally exhibited greater connectivity compared to the unsuccessful group at the pre-quit visit, with these differences normalizing by the post-quit visit (**Figure 1B**). A similar pattern was observed in the SN-ECN connections, where FC in the successful smoking cessation group was higher at the pre-quit visit and stabilized by the post-quit visit (**Figure 1B**). A few exceptions were noted in the SN-ECN, including connections #2, #4, and #6 (refer to **Table 2**), where connectivity in the successful smoking cessation group was similar to the unsuccessful group at the pre-quit visit, but it decreased at the post-quit visit, falling below the unsuccessful group (**Figure 1B**).

### Relationship to Smoking-Related Measures

Correlations were calculated between the FC values from each significant connection and smoking-related variables, resulting in 77 tests (11 FC values x 7 smoking-related variables). Of note, only pre-quit visit correlations were analyzed, as the smoking-related questionnaires were administered only at the pre-quit visit and not at the post-quit visit. A Bonferroni correction was applied to account for multiple comparisons with a significance threshold set at p < 0.05 / 77 (p < 0.0006). After applying this correction, none of the correlations reached statistical significance. However, several trends emerged with uncorrected p < 0.05, suggesting potential relationships that warrant further investigation.

The most notable correlation findings were observed in the SN-ECN connections of the successful smoking cessation group. For the FC strength between the SN (middle frontal gyrus) and ECN (DLPFC) at the pre-quit visit (connection #2), a positive correlation was found with nicotine withdrawal severity, as measured by the CWS (r = 0.4, uncorrected p = 0.04; **Figure 2A**), as well as with smoking duration (r = 0.49, uncorrected p < 0.05). Additionally, for the FC strength between the SN (ACC) and ECN (inferior parietal cortex) at the pre-quit visit (connection #4), a positive correlation with nicotine withdrawal severity was observed in the successful quit group (r = 0.43, uncorrected p = 0.027; **Figure 2B**). In the unsuccessful quit group, weak associations were also found with nicotine dependence severity, as measured by the FTND (r = 0.28, uncorrected p < 0.05) and CDS (r = 0.32, uncorrected p < 0.05) for the same connection. Furthermore, significant positive correlations were identified between three other SN-ECN connections (connections #5, #9, and #10) and smoking duration. The complete correlation results are shown in **Supplementary Figures 2 and 3**.

**Figure 2.**
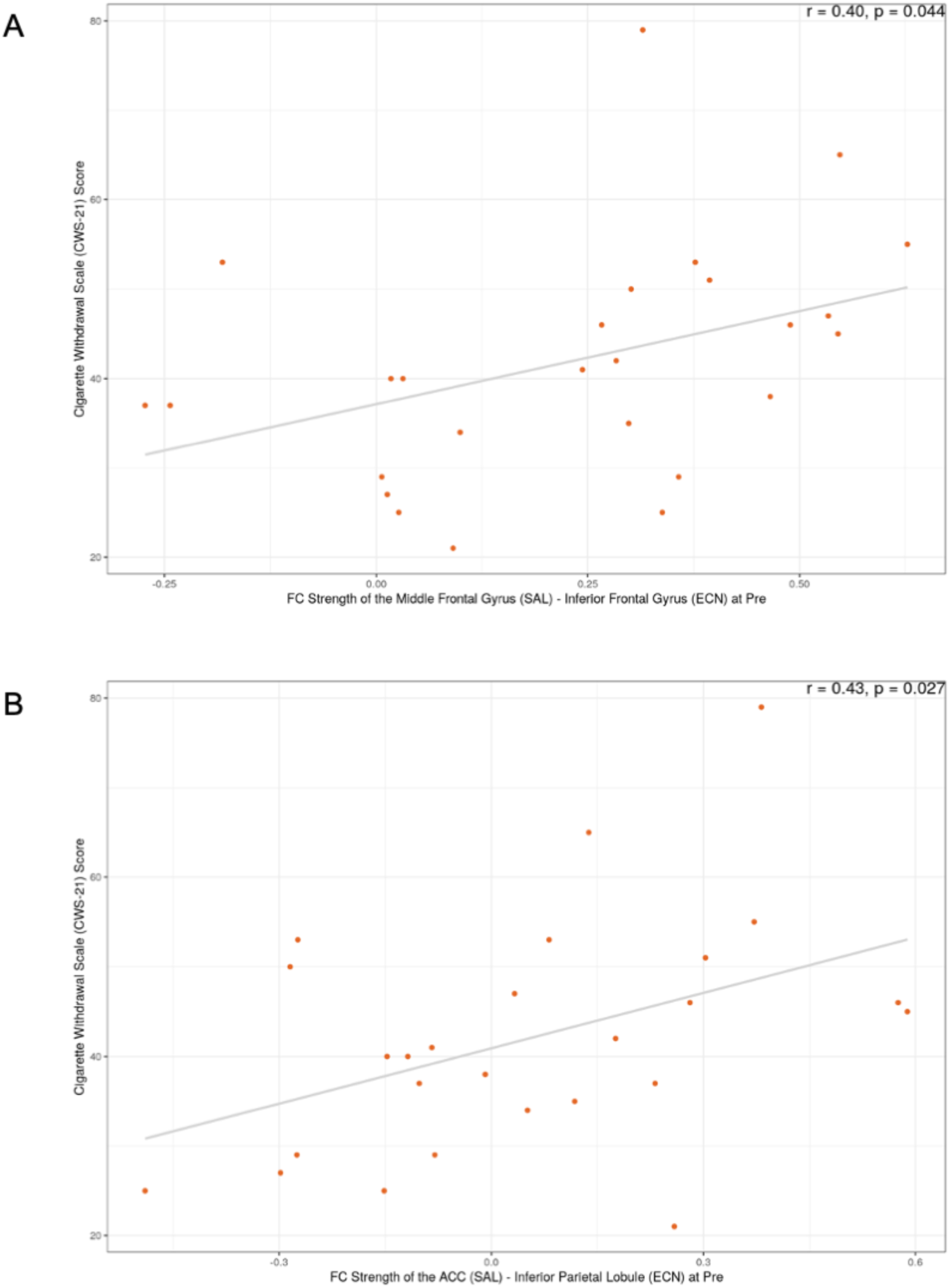
(A) Correlation plot between FC strength of the SN (middle frontal gyrus) – ECN (DLPFC) at the pre-quit visit and the Cigarette Withdrawal Scale (CWS) score in the successful smoking cessation group. (B) Correlation plot between FC strength of the SN (ACC) – ECN (Inferior parietal cortex) at the pre-quit visit and the CWS-21 score in the successful smoking cessation group.

## 4. Discussion

The present study investigated the dynamic changes in FC patterns within and between the large-scale brain networks during a 6-week smoking cessation period, specifically focusing on the SN, ECN, and DMN, and comparing these patterns between successful quitters and unsuccessful quitters. Our network analysis revealed significant group-by-time interaction effects on FC, driven by changes in inter-network connectivity between the SN–DMN and SN–ECN. A key finding was observed in the SN–DMN connections: the successful quit group exhibited relatively higher FC in these pathways at the pre-quit visit compared to the unsuccessful quit group, with these differences largely normalizing by the post-quit visit. Similarly, the SN–ECN connections in the successful quit group showed higher FC at the pre-quit visit relative to the unsuccessful quit group, with most of these connections stabilizing at comparable levels by the post-quit visit. A few exceptions were observed in the SN-ECN connections, where FC, initially comparable between the groups at the pre-quit visit, declined in the successful cessation group at the post-quit visit.

The observed group-by-time interactions offer several interpretative possibilities. The SN is known to mediate the detection of salient stimuli and to facilitate dynamic switching between the internally directed DMN and the externally focused ECN (36, 37). Neuroimaging studies have shown that during cognitively demanding tasks, FC between the SN and ECN is enhanced to allocate attention to external stimuli, while FC between the SN and DMN is decreased to reduce internal self-referential processing (38). In a similar vein, prior neuroimaging research in addiction shows that during withdrawal and craving states, this pattern is disrupted, with a reduction in connectivity between the SN and ECN and increased connectivity between the SN and DMN, reflecting a shift toward internally focused attention and impaired cognitive control (39). This shift is thought to destabilize cognitive control, promote internally directed attention, and exacerbate craving symptoms, thereby increasing relapse vulnerability. Based on this, we hypothesized that successful quitters would show a decrease in SN-DMN connectivity and an increase in SN-ECN connectivity. However, unexpectedly, we observed an increase in both SN-DMN and SN-ECN connectivity at the pre-quit visit, which was normalized by the post-quit visit, with a few exceptions.

Accumulating evidence challenges the traditional view, suggesting that network interactions among the SN, DMN, and ECN dynamically change depending on task goals and demands (40). Notably, some studies have shown that in working memory tasks with increasing cognitive load, the SN exhibits increased connectivity with the DMN, in addition to its enhanced coupling with the ECN (41, 42). These findings imply that the modulation between these networks is not always unidirectional; instead, under certain conditions, there may be an initial increase in SN-DMN connectivity, reflecting an attempt to integrate or monitor internally generated signals. In this context, the higher pre-quit connectivity observed in the successful quit group in both SN-DMN and SN-ECN connections may reflect an adaptive allocation of cognitive resources and interoceptive awareness, enabling individuals to more effectively manage cessation-induced symptoms. Furthermore, the normalization of these connectivity patterns may imply a return to baseline levels following successful cessation.

Our correlational analysis of FC metrics and smoking-related measures may further help interpret the results. Although none of these relationships reached significance after correcting for multiple comparisons, FC between the SN (middle frontal gyrus) and ECN (DLPFC) showed moderate positive correlations with both withdrawal severity and smoking duration in the successful quit group. Similarly, FC between the SN (ACC) and ECN (inferior parietal cortex) was positively related to withdrawal severity in the successful quit group. These findings suggest that higher baseline FC within SN–ECN circuits may be associated with greater withdrawal symptoms and longer smoking histories. Interestingly, these two connections were among the exceptions where baseline FC showed no group differences, but later decreased in successful quitters compared to unsuccessful quitters. One possible interpretation of the decrease is a downregulation of overactive salience signals, which may support the reestablishment of a balanced neural state. This reduction in hypervigilance and excessive attentional bias toward withdrawal-related cues, factors frequently implicated in relapse, could ultimately facilitate successful cessation (43).

Our findings contribute to the growing body of literature suggesting that the functional interplay between large-scale brain networks—particularly among the SN, DMN, and ECN—plays a pivotal role in nicotine addiction and smoking cessation outcomes (9, 15). In successful quitters, the higher pre-quit FC within SN–DMN and SN–ECN connections might reflect an initially heightened state of neural communication geared toward processing interoceptive and salient information. Such a state, however, appears to ‘normalize’ once smoking cessation is achieved, potentially indicating a neural recovery or reorganization process that underpins successful behavior change.

It is also noteworthy that no significant group differences were identified at the individual pre- and post-quit visits; rather, the significant effects emerged only in the group-by-time interaction analysis. This pattern suggests that it is not the absolute level of connectivity at a given moment that matters, but rather the dynamic changes in connectivity over time that are most predictive of cessation outcomes. This dynamic perspective is consistent with recent data-driven connectivity approaches that emphasize the importance of capturing both shared and unique individual-level network alterations over the course of treatment (44).

In summary, our findings suggest that successful smoking cessation is associated with distinct alterations in FC across key brain networks involved in attention, salience, and cognitive control. In particular, a higher pre-quit FC in SN–DMN and SN–ECN connections, which normalizes or is selectively modulated during abstinence, appears to be an important marker of cessation success. Furthermore, the observed trends linking SN–ECN connectivity to nicotine withdrawal severity and smoking duration highlight the potential clinical relevance of these network measures as biomarkers of treatment response. Although these relationships did not survive very conservative statistical correction, they provide a compelling impetus for further investigation. Future studies are needed to replicate and extend these findings, ideally incorporating larger sample sizes, longitudinal designs, and complementary analytic techniques such as dynamic FC and graph theory approaches. Such work could further clarify the role of network-level connectivity alterations as predictors of smoking cessation outcomes and identify novel targets for neuromodulation or pharmacological intervention.

Overall, this study contributes to a more nuanced understanding of the interplay between large-scale brain networks in the context of smoking cessation. By focusing on changes in inter-network connectivity over time, our findings underscore the importance of dynamic neural reorganization during the cessation process. Such reorganization, as reflected in the normalization of SN–DMN and SN–ECN interactions, may represent a key neurobiological mechanism that supports successful abstinence. The identification of these network markers has significant implications for developing targeted, personalized interventions that address both the cognitive and affective dimensions of nicotine withdrawal and addiction.

## Acknowledgments

This research was supported by the Basic Science Research Program through the National Research Foundation (NRF) of Korea funded by the Ministry of Science, ICT, & Future Planning (RS-2018-NR031737, RS-2021-II211343, RS-2024-00435727, RS-2025-00516410) to W.-Y.A.; and the ‘Future Star Psychologist Award’ from the Institute of Psychological Science at Seoul National University to J.J.I..

