## Supplemental File for "Dynamic Brain Network Changes Associated with Successful Smoking Cessation"

**Supplementary Information**

A

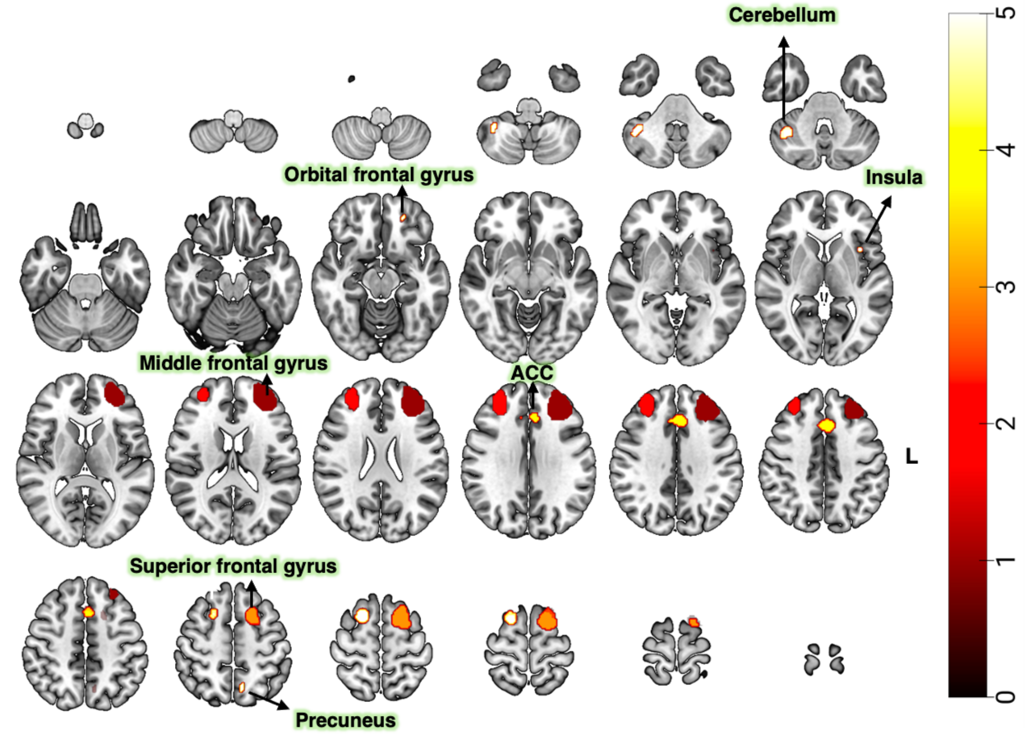


B

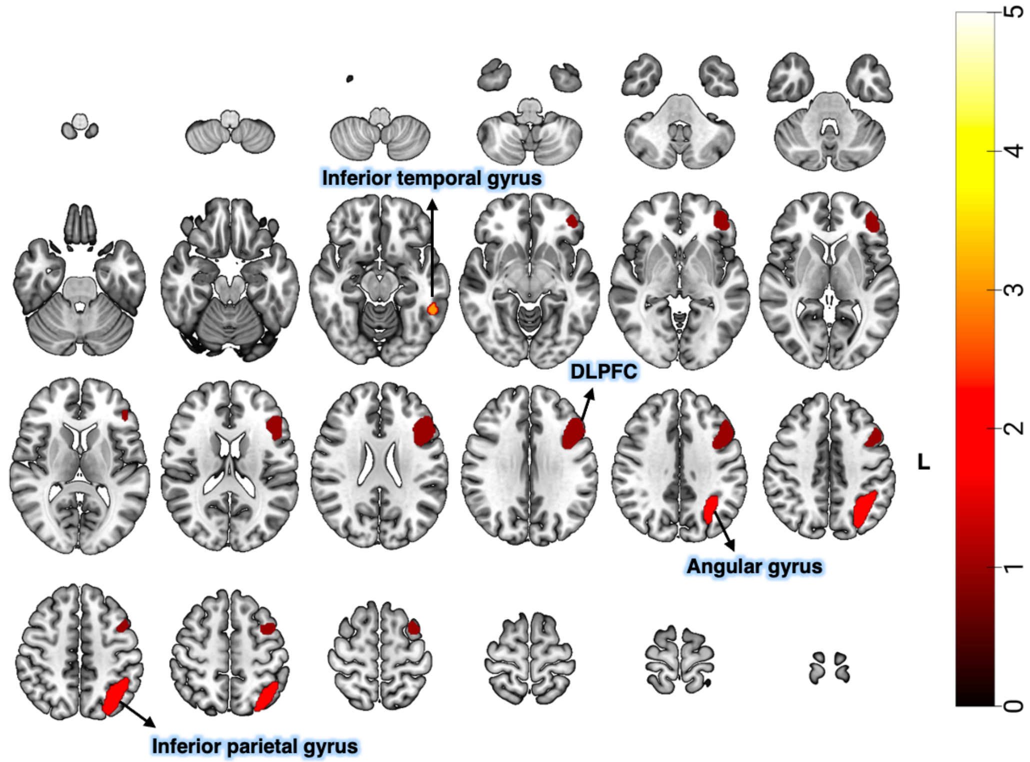

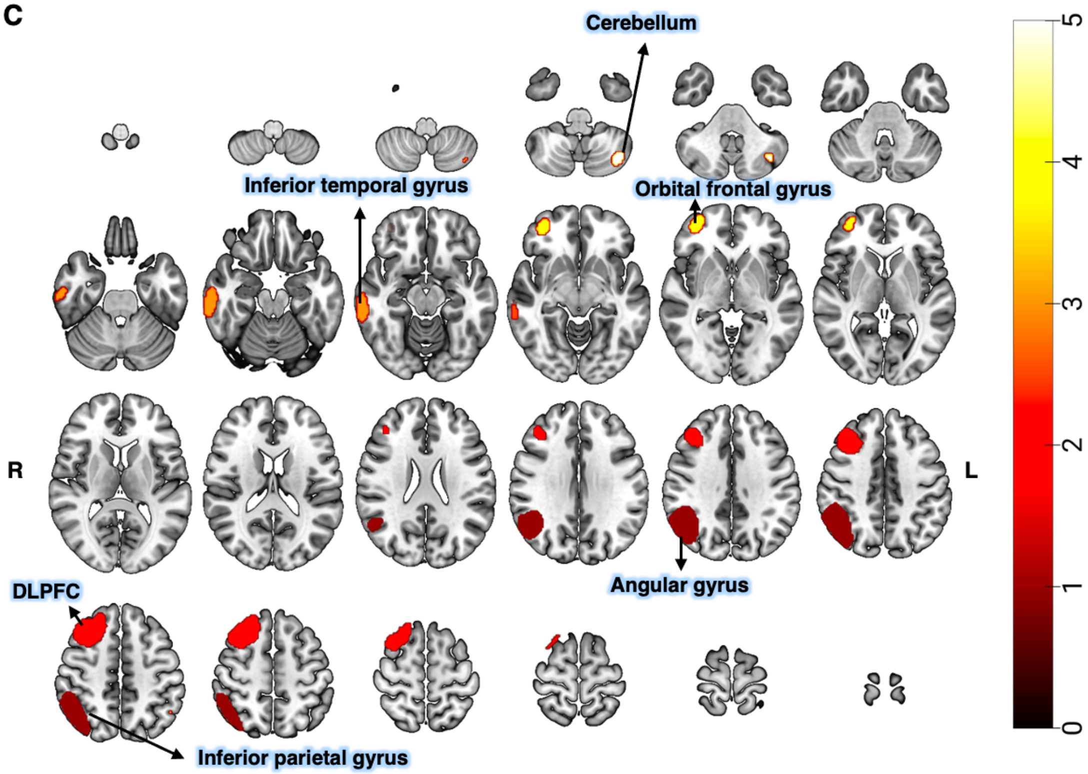
C
Cerebellum
Inferior temporal gyrus
Orbital frontal gyrus
R
L
Angular gyrus
DLPFC
Inferior parietal gyrus
5
4
3
2
1
0

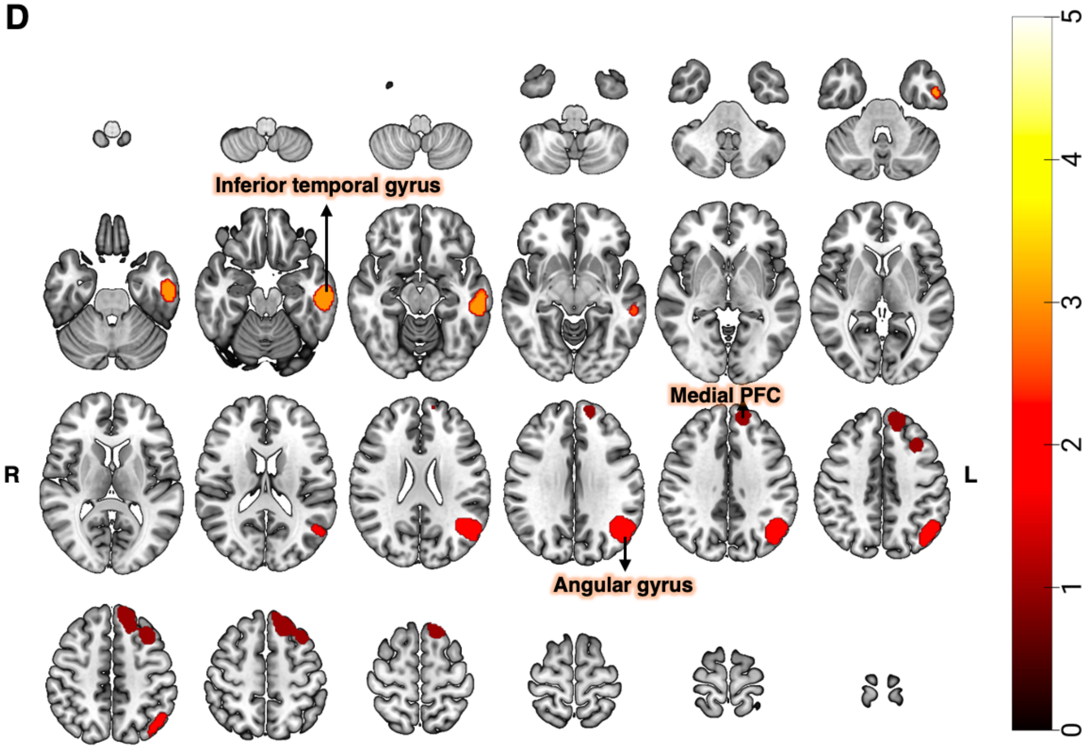
D
Inferior temporal gyrus
Medial PFC
R
L
Angular gyrus
5
4
3
2
1
0

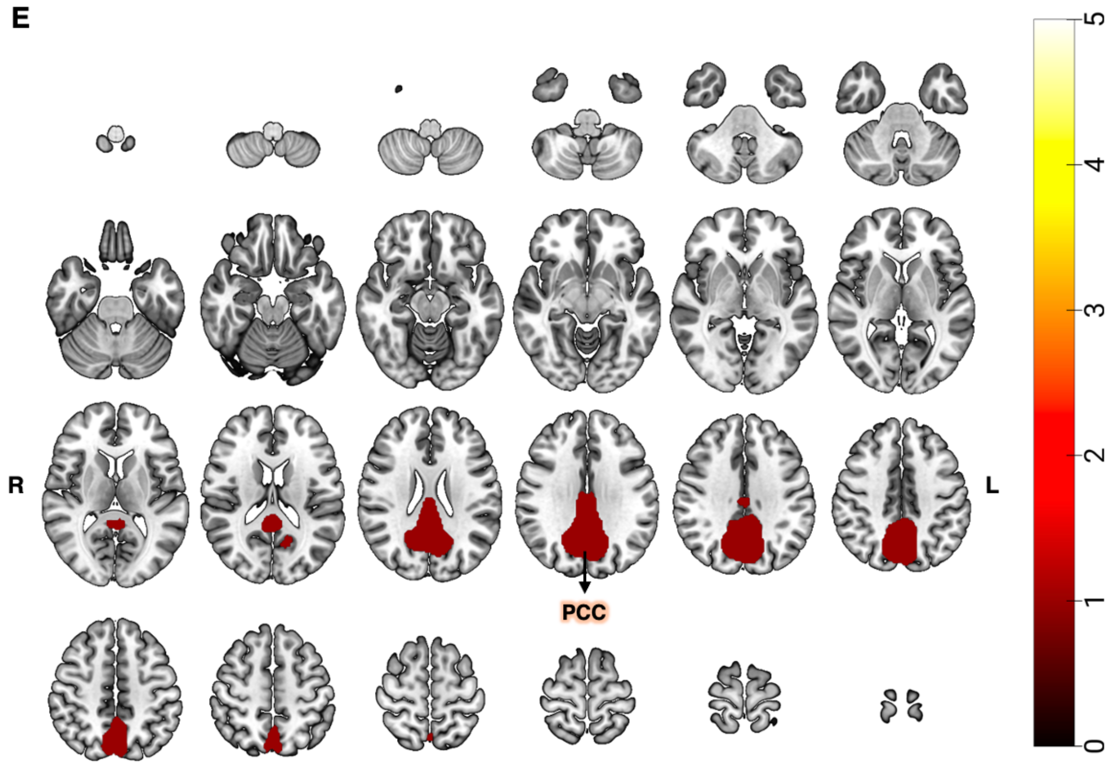


**Supplementary Figure 1.** ICA results. (A) Salience network, (B) Left ECN, (C) Right ECN, (D) DMN, (E) DMN.

Abbreviations: ACC: Anterior Cingulate Cortex; DLPFC: Dorsolateral Prefrontal Cortex; PCC: Posterior Cingulate Cortex; PFC: Prefrontal Cortex.

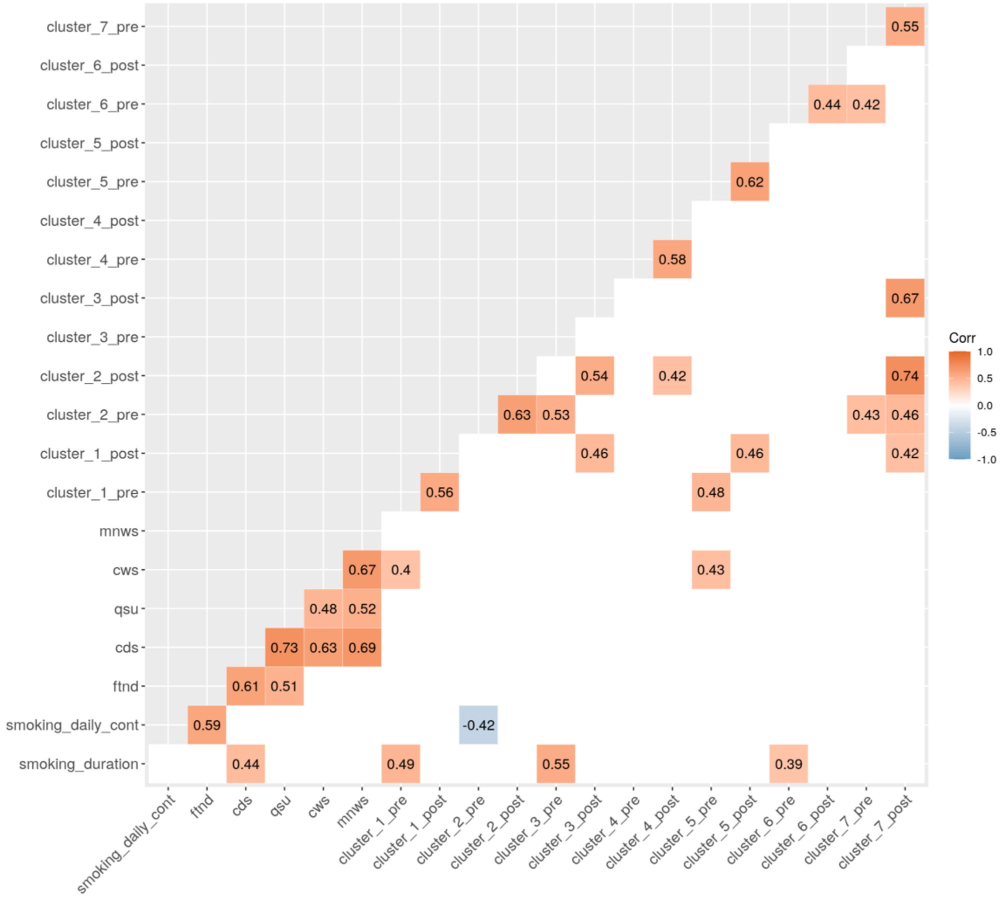


**Supplementary Figure 2.** Correlation plot of functional connectivity values from the significant clusters and smoking-related variables for the successful quit group.

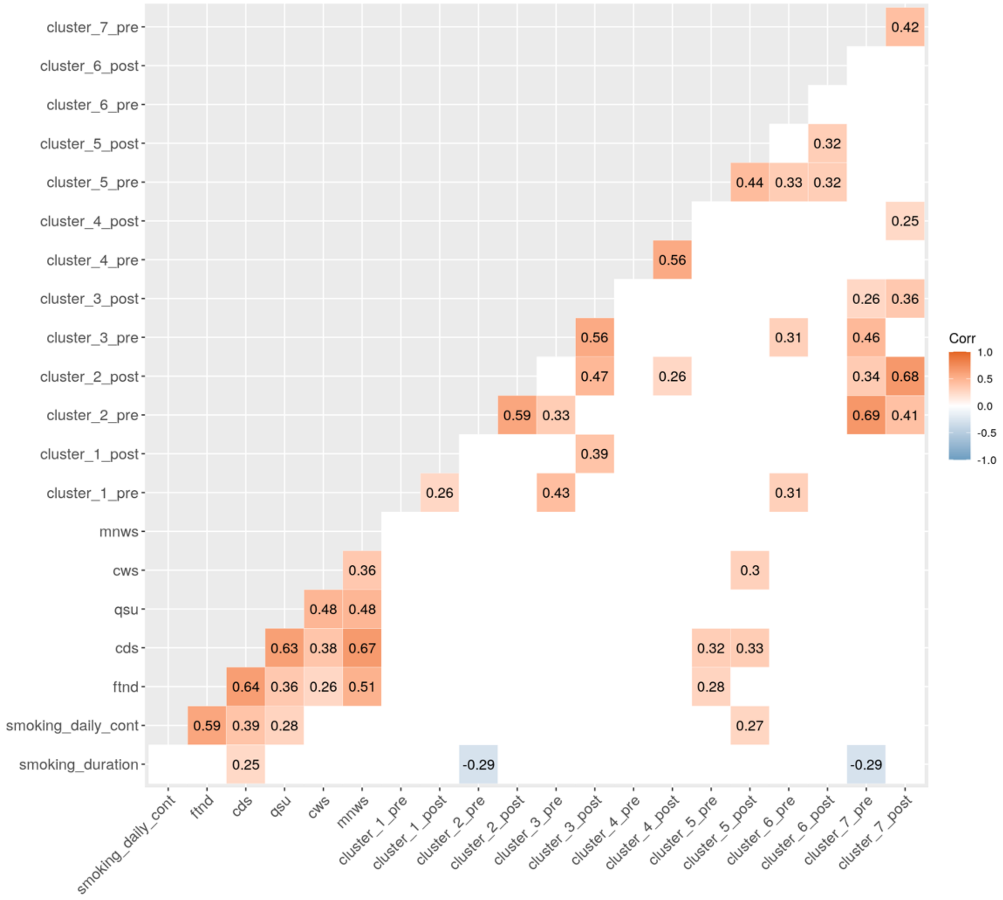


**Supplementary Figure 3**. Correlation plot of functional connectivity values from the significant clusters and smoking-related variables for the unsuccessful quit group.
